# Loss of the predicted cell adhesion molecule MPZL3 promotes EMT and chemoresistance in ovarian cancer

**DOI:** 10.1101/2024.11.14.623672

**Authors:** Ya-Yun Cheng, Beth L. Worley, Zaineb Javed, Amal T. Elhaw, Priscilla W. Tang, Sarah Al-Saad, Shriya Kamlapurkar, Sierra White, Apoorva Uboveja, Karthikeyan Mythreye, Katherine M. Aird, Traci A. Czyzyk, Nadine Hempel

**Affiliations:** Department of Medicine, Division of Malignant Hematology & Medical Oncology, UPMC Hillman Cancer Center, University of Pittsburgh, PA, USA; Department of Pharmacology, College of Medicine, Pennsylvania State University, Hershey, PA, USA; Department of Pharmacology & Chemical Biology, UPMC Hillman Cancer Center, University of Pittsburgh School of Medicine, Pittsburgh, PA; Department of Pathology and O’Neal Comprehensive Cancer Center, Heersink School of Medicine, University of Alabama at Birmingham, Birmingham, AL, USA; Department of Anesthesiology & Perioperative Medicine, Penn State University College of Medicine, Hershey, PA, USA

**Author notes:** Corresponding Author: Nadine Hempel, PhD, University of Pittsburgh School of Medicine UPMC Hillman Cancer Center, The Assembly, Rm 2039, 5051 Centre Ave, Pittsburgh, PA 15213. Current: Department of Cardiometabolic Disease, Merck Research Laboratories., South San Francisco, CA, USA. The authors declare no potential conflicts of interest.

## Abstract

Myelin protein zero-like 3 (MPZL3) is an Immunoglobulin-containing transmembrane protein with predicted cell adhesion molecule function. Loss of 11q23, where the *MPZL3* gene resides, is frequently observed in cancer, and *MPZL3* copy number alterations are frequently detected in tumor specimens. Yet the role and consequences of altered MPZL3 expression have not been explored in tumor development and progression. We addressed this in ovarian cancer, where both MPZL3 amplification and deletions are observed in respective subsets of high-grade serous specimens. While high and low MPZL3 expressing populations were similarly observed in primary ovarian tumors from an independent patient cohort, metastatic omental tumors largely displayed decreased MPZL3 expression, suggesting that MPZL3 loss is associated with metastatic progression. MPZL3 knock-down leads to strong upregulation of vimentin and an EMT gene signature that is associated with poor patient outcomes. Moreover, MPZL3 is necessary for homotypic cancer cell adhesion, and decreasing MPZL3 expression enhances invasion and clearance of mesothelial cell monolayers. In addition, MPZL3 loss abrogated cell cycle progression and proliferation. This was associated with increased resistance to Cisplatin and Olaparib and reduced DNA damage and apoptosis in response to these agents. Enhanced Cisplatin resistance was further validated *in vivo*. These data demonstrate for the first time that MPZL3 acts as an adhesion molecule and that MPZL3 loss results in EMT, decreased proliferation, and drug resistance in ovarian cancer. Our study suggests that decreased MPZL3 expression is a phenotype of ovarian cancer tumor progression and metastasis and may contribute to treatment failure in advanced-stage patients.

## Introduction

Ovarian cancer is the second most lethal gynecological malignancy in the United States, responsible for over 12,000 deaths annually and with a 5-year overall survival rate of less than 27% for patients with stage IV high-grade serous adenocarcinoma, which is the most common ovarian cancer histologic subtype [1, 2]. More than 60% of patients are diagnosed after stage III, due in part to both lack of effective screening methods and asymptomatic disease progression until metastasis to the peritoneal cavity [1]. Ovarian cancer frequently undergoes metastasis through transcoelomic pathways. This process involves the detachment of cells from the primary tumor site, leading to their spread as either individual cells or spheroids into the peritoneal cavity, a process that is significantly influenced by alterations in cell-cell and cell-extracellular matrix (ECM) adhesion [3–7]. Depending on the type of cell adhesion molecule (CAM), both loss and gain of CAMs can contribute to metastatic progression.

Myelin protein zero-like 3 (MPZL3) is an Immunoglobulin (Ig) domain-containing transmembrane protein, that shares sequence homology with junctional adhesion molecule family members, including V-set and immunoglobulin domain containing 1 (VSIG1). The function of MPZL3 has primarily been investigated through studies in murine knock-out models, where it has been associated with epidermal barrier formation, sebaceous gland differentiation, hair follicle cycling, and metabolic regulation [8–15]. In addition, we have shown that even transient knock-down of MPZL3 can prevent the negative metabolic effects of a high-fat diet in mice [8]. Furthermore, MPZL3 can also localize to mitochondria [16]. However, it remains largely unclear if and how these functions relate to MPZL3’s role as a CAM.

Junctional adhesion molecules that share sequence homology with MPZL3 and other MPZ myelin zero-like proteins are often highly expressed in normal polarized epithelia and endothelial cells. The loss of VSIG1 was shown to be a feature of gastric, lung, and esophageal cancer cells and necessary for metastasis [17]. Whether MPZL3 has similar properties in tumor cells has not been explored [10]. Interestingly, the *MPZL3* locus (11q23.3) is susceptible to chromosomal loss in multiple cancer subtypes [18–21], and loss of heterozygosity of this location is a frequent event in ovarian cancer [19]. This suggests that loss of MPZL3 may be a phenotype of cancer. Previous research indicates varying MPZL3 expression patterns across different cancer types, suggesting that the role of MPZL3 is context-dependent [22]. High MPZL3 expression has been associated with poor prognosis in breast cancer and shown to drive the proliferation of MET-amplified cancer cell lines [22, 23].

Here we set out to explore the role of MPZL3 in ovarian cancer and found that loss of MPZL3 results in loss of homotypic cell adhesion and a more invasive phenotype associated with epithelial to mesenchymal transition (EMT). In addition, MPZL3 knock-down slows cell cycle progression, promotes a senescent-like phenotype, and contributes to chemoresistance *in vitro* and *in vivo*. These data demonstrate that decreased MPZL3 expression is a phenotype of ovarian cancer tumor progression and metastasis and may contribute to treatment failure in advanced stage disease patients.

## Materials and methods

### Cell culture

All cell lines were cultured in a 5% CO_2_ environment at 37 °C. OVCAR4 cells (MilliporeSigma, SCC258) were cultured in RPMI-1640 (Gibco, 11875) supplemented with 2 mM glutamine (Gibco, 25030081), 10% FBS (Avantor Seradigm, 1500-500), 0.25 U/mL insulin (Gibco, 12585014). OVCA433 was kindly provided by Dr. Susan K. Murphy (Duke University) and cultured in RPMI 1640 medium (Corning,10-040-CV) supplemented with 10% FBS. SKOV3-Luc with stably expressed Luciferase was a gift from Dr. Karthikeyan Mythreye (the University of Alabama at Birmingham) and maintained in RPMI 1640 medium (Corning,10-040-CV) supplemented with 10% FBS. 293FT was purchased from Invitrogen (R70007) and cultured in DMEM (Corning, 10- 017-CV) with 10% FBS. NIH:OVCAR3 (OVCAR3) and OV90 cells were purchased from the American Type Culture Collection (ATCC, HTB-161, CRL-3585). OVCAR3 was maintained in RPMI 1640 medium (Gibco, 11875) supplemented with 10% FBS and 0.01 mg/ml bovine insulin (MilliporeSigma, I0516). OV90 was maintained in a 1:1 mixture of MCDB 105 medium (Cell Applications, 117-500) with 1.5 g/L sodium bicarbonate (Gibco, 25080094) and Medium 199 (Corning,10-060-CV) with 2.2 g/L sodium bicarbonate, supplemented with 15% FBS. FT282 was kindly provided by Dr. Ronny Drapkin (University of Pennsylvania) and cultured in 50% DMEM and 50% Ham’s F-12 medium (Corning,10-090-CV) supplemented with 2% FBS. GFP-labeled mesothelial cells, ZTGFP, was a gift from Dr. Ioannis Zervantonakis (University of Pittsburgh) [24] and maintained in 1:1 mixture of Medium 199 (Corning,10-060-CV) and MCDB105 (Cell Applications, 117-500) media, supplemented with 10% FBS FBS and 1% Penicillin-Streptomycin (GIBCO). Cells are routinely tested for mycoplasma with EZ-PCR Mycoplasma Detection Kit (Captivate Bio,20-700-20) and authenticated using STR sequencing (Labcorp).

### Cloning and lentiviral transduction

MPZL3 shRNAs were from TRC library (MilliporeSigma), with the following targeting sequences. shMPZL3 #1: GAGTCACCTAAAGACAGGAAA (TRCN0000137304). shMPZL3 #2: GGGATGCATCTATAAGTATAA (TRCN0000413087). shMPZL3 #3: GCAGCCACACAGTATCAATAT (TRCN0000137252). pLKO.1 scramble (scr) shRNA and pLKO.1 control shRNA (Addgene, #8453) were used as controls. The expression or shRNA plasmids were co-transfected with the virus package plasmids, psPAX2 (Addgene, #12260) and pMD2.G (Addgene, #12259), to 293FT cells for virus production. Cells were seeded on 6-well plates the day before virus transduction. 200 µL of virus was added to the cells with polybrene (MilliporeSigma, TR-1003) (0.8 µg/mL) and incubated for 24 hrs. Drug selection was performed 3 days after transduction, with the concentrations listed below. OVCA433: 2 µg/mL of puromycin (Gibco, A1113803); SKOV3-Luc: 3 µg/mL of puromycin; OVCAR4: 2 µg/mL of puromycin.

### RNA isolation and semi-quantitative real-time RT-PCR

Total RNA was isolated by Direct-zol RNA Miniprep kit (Zymo Research, R2052) and used for cDNA synthesis (Quantabio: 95047), based on the manufacturer’s instruction. cDNA samples were analyzed by semi-quantitative real time RT-PCR using Bio-Rad CFX Opus 96 Real-Time PCR System, with PowerUp SYBR Green Master Mix (Applied Biosystems) or iTaq Universal SYBR Green Supermix (Bio-Rad). Primer sequences used: MPZL3 sense: 5’- ATGCCCATGTCCGAGGTTATG -3’, MPZL3 antisense: 5’- GGAGGGCGATATGTCCAGTCT-3’; GAPDH sense: 5’-GAGTCAACGGATTTGGTCGT-3’, GAPDH antisense: 5’- TTGATTTTGGAGGGATCTCG -3’; ACTB sense: 5’- AGAGCTACGAGCTGCCTGAC-3’; ACTB antisense: 5’- AGCACTGTGTTGGCGTACAG-3’. RNA relative expression was normalized to the reference gene (GAPDH and ACTB) using the 2^-ΔΔCT^ formula, and relative changes between samples were calculated by normalizing to scr control cells.

### Patient data analysis

MPZL3 copy number alteration (CNA) incidence was examined on cBioportal with the following datasets: pan-cancer analysis of whole genomes (ICGC/TCGA) [25] and ovarian serous cystadenocarcinoma (TCGA, PanCancer Atlas). Overall survival and transcriptome data for ovarian serous cystadenocarcinoma (TCGA, PanCancer Atlas) dataset were obtained from cBioportal. For analyzing transcriptome changes according to the expression of MPZL3, the data were divided into quantiles based on the expression level of MPZL3 RNA. By comparing the first (low MPZL3) and the fourth (high MPZL3) quantiles, the genes that showed significantly higher expression in each group were selected for further analysis. Volcano plots were generated using VolcaNoseR software. Pathway analysis was performed using Molecular Signatures Database (MSigDB) analysis to compute overlaps between the selected genes and the gene sets in MSigDB [26–28]. Here, we applied Hallmark gene set for analysis [28].

### Tumor specimens

Specimens of normal fallopian tube, ovarian tumor and omental tumor from high grade serous ovarian cancer patients were obtained through an honest broker from the ProMark biospecimens bank at the University of Pittsburgh, with IRB approval from the University of Pittsburgh (Protocol # STUDY21050194). RNA was isolated and RT-PCR was carried out as above.

### RNA Sequencing

OVCAR4 cells were transduced with scramble or MPZL3 shRNA (sh #1 and sh #2) virus and selected with puromycin for 3 days. OVCA433 cells were expressing empty vector control or MPZL3 shRNA (sh #1 and sh #3). Total RNA was extracted from cells using Direct-zol RNA Miniprep kit (Zymo Research, R2052). The samples were sequenced using the Illumina NovaSeq 6000 platform and sequence alignment was conducted with Hisat2 mapping tool. Differential expression analysis was performed with DESeq2 R package. The adjusted P-value ≤0.05 was used to define differentially expressed genes. The data generated in this study are publicly available in Gene Expression Omnibus (GEO; GEO GSE accession pending)

### Protein isolation and western blotting

Cells were scraped and harvested in RIPA buffer (Thermo Scientific, 89901) with protease and phosphatase inhibitors (Thermo Scientific, 78443). Protein concentration was measured with the Bradford protein assay (Bio-Rad, 5000006). 30 µg of total protein was loaded to 4–20% SDS-PAGE gels (Bio-Rad) and transferred to PVDF membranes (Thermo Scientific, PI88520) after electrophoresis. The membranes were blocked with 5% non-fat milk (Bio-Rad,1706404) in TBS containing 0.1% Tween20 (MilliporeSigma, 900-64-5) for 30 mins, and incubated with primary antibodies at 4°C, overnight. The primary antibodies used in this study included MPZL3 (Novus Biologicals, NBP2-84169, 1:500), N-cadherin (Cell Signaling, #14215, 1:1000), Vimentin (Cell Signaling, #5741, 1:1000), and GAPDH (Santa Cruz, #sc-47724, 1:1000). The next day, after washing with 0.1% TBST, the membranes were incubated with HRP-conjugated secondary antibodies (Cytiva, NA931 and NA934, 1:10000) at room temperature for 1 hr. After washing, the signals were detected using SuperSignal West Femto Maximum Sensitivity Substrate (Thermo Scientifi, 34096) and ChemiDoc XRS+ system (Bio-Rad).

### Adhesion assay

Parental OVCAR4 or ZTGFP cells were seeded in 96-well plates. The following day, OVCAR4 cells expressing scramble control or MPZL3 shRNA (sh #1 and sh #2) were stained with CellTrace Far Red (Invitrogen, C34572) and added as a suspension to the pre-seeded 96-well plates. After incubating for 30 minutes at 37°C, non-adherent cells were washed off with PBS three times, each for 5 minutes. The adherent cells were then examined under a microscope (Leica Thunder Imager) and analyzed using FIJI software (NIH).

### Spheroid formation and mesothelial clearance assay

OVCAR4 cells expressing scramble control or MPZL3 shRNAs (sh #1 and sh #2) were stained with CellTrace Far Red (Invitrogen, C34572), seeded into 96-well ULA plates (Corning, 7007), and incubated overnight to form spheroids. The following day, the spheroids were transferred to a pre-seeded 96-well plate containing a layer of ZTGFP cells. After 48 hrs of incubation, images were captured using a Leica Thunder Imager and analyzed with FIJI software (NIH) to assess the areas cleared by the spheroids.

### Growth assay and IC_50_ analysis

Cells were seeded into 96-well plates in 100 µL medium with the following conditions. OVCA433: 500 cells/well, OVCAR4: 1000 cells/well, SKOV3-Luc: 500 cells/well. Cell growth assay was performed with FluoReporter Blue Fluorometric dsDNA Quantitation Kit (Invitrogen, F2962). One 96-well plate was harvested, with medium removed, and stored at -80 °C freezer every 24 hrs. Total 5 plates were collected and proceeded to Hoechst 33258 staining, according to the manufacturer’s instruction. The DNA quantification was measured by a Fluorescence plate reader (PerkinElmer), with excitation at 360 nm and detection at 460 nm. The cell growth rate was calculated by normalizing the values to day 1. IC_50_ analysis was performed using OVCAR4 cells with or without MPZL3 knockdown. The cells were seeded and treated with Paclitaxel (MilliporeSigma), Cisplatin (MilliporeSigma), or Olaparib (Selleck Chemicals, S1060) the next day and then incubated for 72 hrs. The plates were harvested and measured using FluoReporter Blue Fluorometric dsDNA Quantitation Kit. IC_50_ values were calculated based on the percentage survival relative to untreated cells.

### Cell cycle analysis

OVCAR4 cells were seeded in 6-well plates at a density of 125000 cells per well and incubated overnight. The following day, the cells were subjected to a double thymidine block for G1/S phase synchronization. Briefly, cells were treated with 2 mM Thymidine (Sigma-Aldrich, T9250) for 18 hrs, released for 8 hours, and then incubated again with 2 mM Thymidine for an additional 16 hrs. Afterward, the cells were placed in fresh media and harvested at 0, 3, 6, 9, and 24 hrs. Following staining with Propidium Iodide (Abcam, ab139418), cell cycle status was assessed by flow cytometry (Beckman Coulter, CytoFLEX) and analyzed using FlowJo software (BD Life Sciences).

### Caspase 3/7 assay

OVCAR4 cells were seeded in 96-well plates at a density of 2000 cells per well. The following day, cells were treated with 1.3 µM Cisplatin (cis-diamineplatinum(II) dichloride) (Sigma) or 3.9 µM Olaparib (Selleck Chemicals, S1060) for 72 hrs. Caspase 3/7 activity was then assessed using the Caspase-Glo 3/7 Assay (Promega, G8091) according to the manufacturer’s protocol. Cells were incubated with an equal volume of lysis reagent for 1 hr at room temperature, protected from light. The samples were transferred to white 96-well plates (BrandTech, BRA-781605), and bioluminescence signals were measured using a GloMax Explorer plate reader (Promega).

### Senescence-associated beta-galactosidase activity assay

OVCAR4 or OVCA433 cells were seeded in 12-well plates at a density of 75000 or 50000 cells per well, respectively, and incubated for 48 hrs. The cells were then fixed with a solution containing 0.2% glutaraldehyde (Polysciences, 01909-100) and 2% formaldehyde (VWR, 0493- 500) for 5 mins. Afterward, a β-Galactosidase staining solution containing 150 mM NaCl (MilliporeSigma), 2 mM MgCl₂ (MilliporeSigma), 5 mM K₃Fe(CN)₆ (MilliporeSigma), 5 mM K₄Fe(CN)₆ (MilliporeSigma), 40 mM Na₂HPO₄ (MilliporeSigma) (pH 5.4 for OVCAR4, pH 5 for OVCA433), and 20 mg/mL X-gal (MilliporeSigma, B4252) was added to the cells. The cells were incubated in a non-CO₂ incubator at 37°C for 16 hrs, then washed three times with ddH₂O and stored in 50% glycerol. Images were taken using a microscope (Invitrogen, EVOS XL Core) at 10x magnification and analyzed with FIJI software (NIH).

### Immunofluorescence (IF)

OVCAR4 cells were seeded on 8-well chambered cell culture slides (Falcon, 08-774-26) for overnight incubation. The next day, the cells were treated with 5 µM Cisplatin (cis-diamineplatinum(II) dichloride) (Sigma) or 3.9 µM Olaparib (Selleck Chemicals, S1060) for 72 hrs. The cells were fixed with 4% PFA (BTC BeanTown Chemical,30525-89-4) for 10 mins, permeabilized with 0.25% Triton X-100 (Fisher Scientific, BP151500) in PBS (Corning, 21-040- CV) for 10 mins, and incubated with blocking buffer containing 2% goat serum/PBS (Cell Signaling Technology, 5425) for 30 mins at room temperature. Then, the cells were incubated with anti-phospho-Histone H2A.X (Ser139) antibody (MilliporeSigma, 05-636, 1:350) in blocking buffer at 4°C overnight. The next day, after washing with PBS three times, the blocking buffer with goat anti-rabbit Alexa Fluor 488 conjugated (Life Technologies, Cat. No. A-11008, 1:1000) antibody was added, incubating at room temperature for 1 hr protected from light. After washing, the cover slips were mounted with ProLong Gold Antifade Reagent with DAPI (Cell Signaling Technology, 8961) and dried at room temperature overnight protected from light. The signals were examined by Leica Thunder Imager at 20x magnification and analyzed by FIJI software (NIH).

### Tumor xenografts

Approval for animal studies was sought from the University of Pittsburgh IACUC prior to study commencement (approved protocol IS00023174). All animals were housed in barrier facilities for immunodeficient mice in the University of Pittsburgh Animal Faculty, and mouse husbandry and experiments were performed in accordance with the guidelines of the Laboratory Animal Ethics Committee of the University of Pittsburgh. OVCAR4 cells expressing scramble or MPZL3 shRNA (sh #1) were mixed with Matrigel (Corning, 354248) for both right and left subcutaneous (flank) injections into female Nod scid gamma mice (NSG, Jackson laboratory; scr: n = 10, sh #1: n = 10, 5 × 10^6^ cells per injection). Mice were randomized prior to drug treatment. Subcutaneous tumor growth was monitored using caliper measurements twice weekly, and tumor volumes were calculated according to the formula V = ½ (length (longer diameter) × width (shorter diameter)^2^). When the average tumor volume in each group reached 50 mm³, mice were injected intraperitoneally (IP) with Cisplatin or vehicle control (saline) twice per week. The Cisplatin dosing regimen consisted of 2 mg/kg body weight for 5 doses, followed by 3 mg/kg body weight for 2 doses, a pause for 3 doses, and then 2 mg/kg body weight for the final 2 doses. One mouse was removed from the study prior to drug treatment due to illness unrelated to tumor burden. One mouse was dead due to Cisplatin treatment, and thus was excluded from the final analysis. The mice were euthanized after completing the treatment regimen (a total of 12 doses, including 3 pauses), and the xenografted tumors were resected and measured. To determine chemotherapy response, the tumor growth rates were calculated by dividing the final tumor volumes by the average tumor volumes from three measures: three days before treatment, on the day treatment began, and four days after treatment began.

### Statistical analysis

Unless otherwise noted, the data presented in this study are based on at least three independent experiments, and are expressed as mean ± SEM. All statistical analyses were performed using GraphPad Prism 10 (GraphPad Software, San Diego, CA), and were selected based on the experimental design, as indicated. Statistical significance was defined as p < 0.05.

## Results

### *MPZL3* gene displays copy number alterations in cancer and MPZL3 expression is decreased in omental metastatic lesions

The *MPZL3* gene is located on chromosome 11q23.3, a region known to be susceptible to chromosomal loss across multiple cancer subtypes [18–20]. Loss of heterozygosity at 11q23 has been observed in breast, lung, colorectal, and cervical cancers [21, 29–34], and is a frequent event in ovarian carcinoma [19]. CNA analysis of ICGC/TCGA Pan-Caner data [25] across 24 tumor types found that many tumor samples exhibit *MPZL3* copy number loss (Figure 1A). Interestingly, esophageal, lung, and ovarian tumors display both loss and gain of *MPZL3.* Approximately 30% of ovarian adenocarcinoma samples have *MPZL3* heterozygous loss, while 25% exhibit chromosomal gain (Supplementary Figure S1A), and these CNAs correlate with MPZL3 mRNA transcript levels (Figure 1B). In addition, MPZL3 transcript expression was negatively associated with gene methylation status (Figure 1C). Analysis of patient survival demonstrated that *MPZL3* copy number loss was associated with a 9.2 month decrease in overall patient survival, although this association was not statistically significant (P=0.2621) (Supplementary Figure S1B). To further assess the expression of MPZL3 during tumor progression, we determined MPZL3 transcript levels in primary ovarian tumors and omental metastatic lesions relative to expression levels in normal fallopian tube specimens from an independent tissue bank at the University of Pittsburgh. Similar to TCGA data, primary tumors were comprised of both high and low-expressing populations, while the majority of omental tumors displayed decreased MPZL3 expression (Figure 1D), suggesting a loss of MPZL3 is associated with tumor progression.

**Figure 1.**
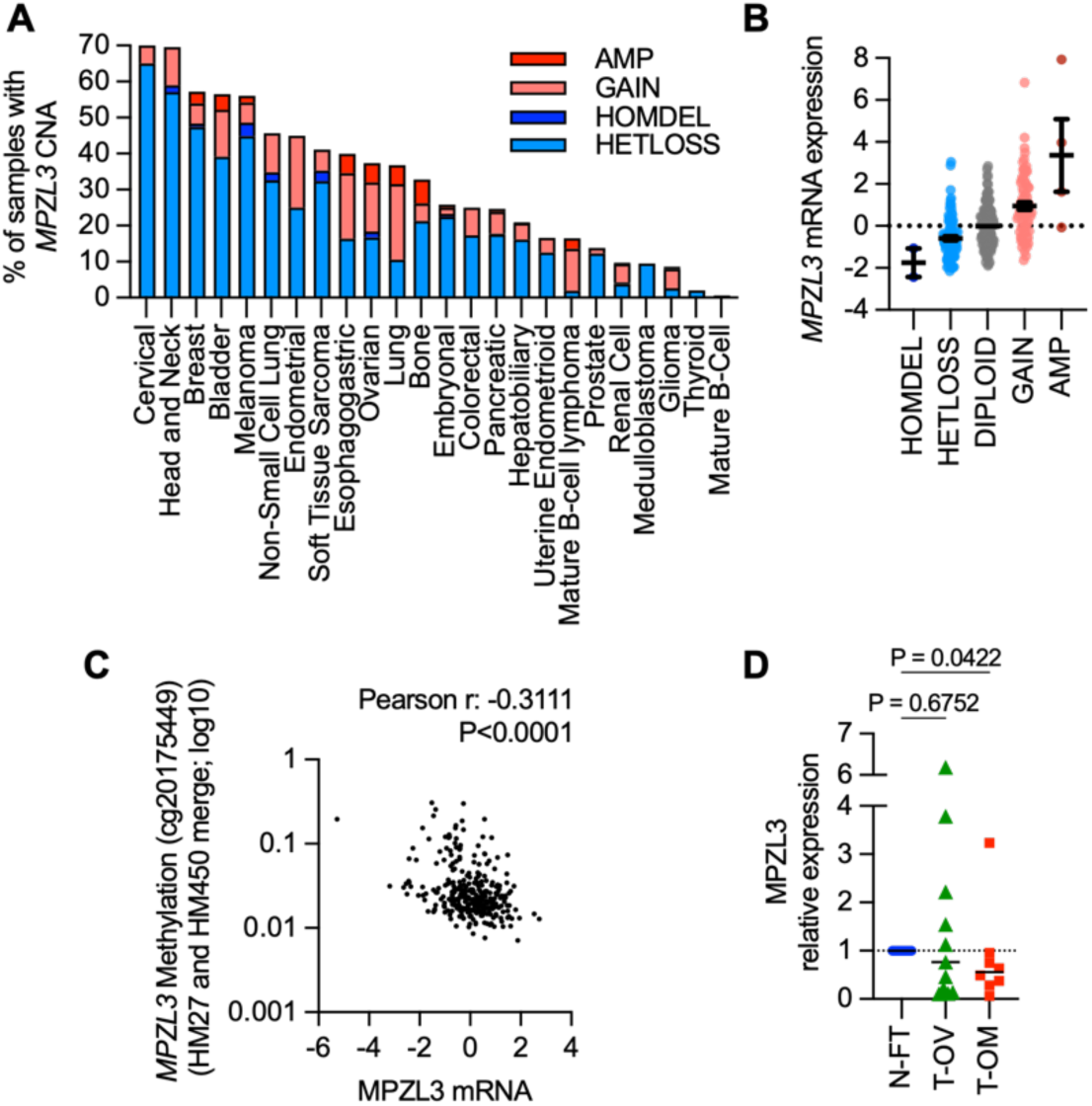
MPZL3 expression in ovarian cancer tissues. A. Percentage of tumor samples displaying *MPZL3* CNA based on ICGC/TCGA pan-cancer analysis of whole genomes. B. MPZL3 mRNA expression and *MPZL3* CNA in ovarian adenocarcinoma samples (TCGA, PanCancer). C. MPZL3 methylation is negatively associated with MPZL3 mRNA expression (TCGA, Pan-Cancer). D. MPZL3 transcript levels detected by sq-RT-PCR in high-grade serous ovarian (T-OV, n=11) and omental tumors (T-OM, n=8) compared to normal fallopian tube tissues (N-FT, n=14; Kruskal-Wallis multiple comparison test P=0.07; Dunn’s multiple comparison test P values shown).

### EMT gene expression is significantly enriched following MPZL3 knockdown

To better understand the role of MPZL3 in ovarian cancer, RNA sequencing was carried out following MPZL3 knock-down in two cell lines of high-grade serous origin, using two independent shRNAs each (Figure 2A-C, Supplementary Figure S2B). OVCAR4 and OVCA433 cells were chosen as these express median levels of MPZL3 compared to other ovarian cancer cell lines (Supplementary Figure S2A). MSigDB analysis was carried out on differentially expressed genes (DEGs, adjusted p-value <0.05 and log2 fold-change ≥1) that were significantly altered by both shRNAs and shared by OVCAR4 and OVCA433 cells following MPZL3 knock-down (Figure 2B&C). Hallmark analysis revealed the strongest enrichment in EMT-related genes following MPZL3 knock-down, while cell cycle-associated gene sets including MTORC1, E2F and G2M checkpoint were enriched in the control group (Figure 2D, Supplementary Figure S2C).

**Figure 2.**
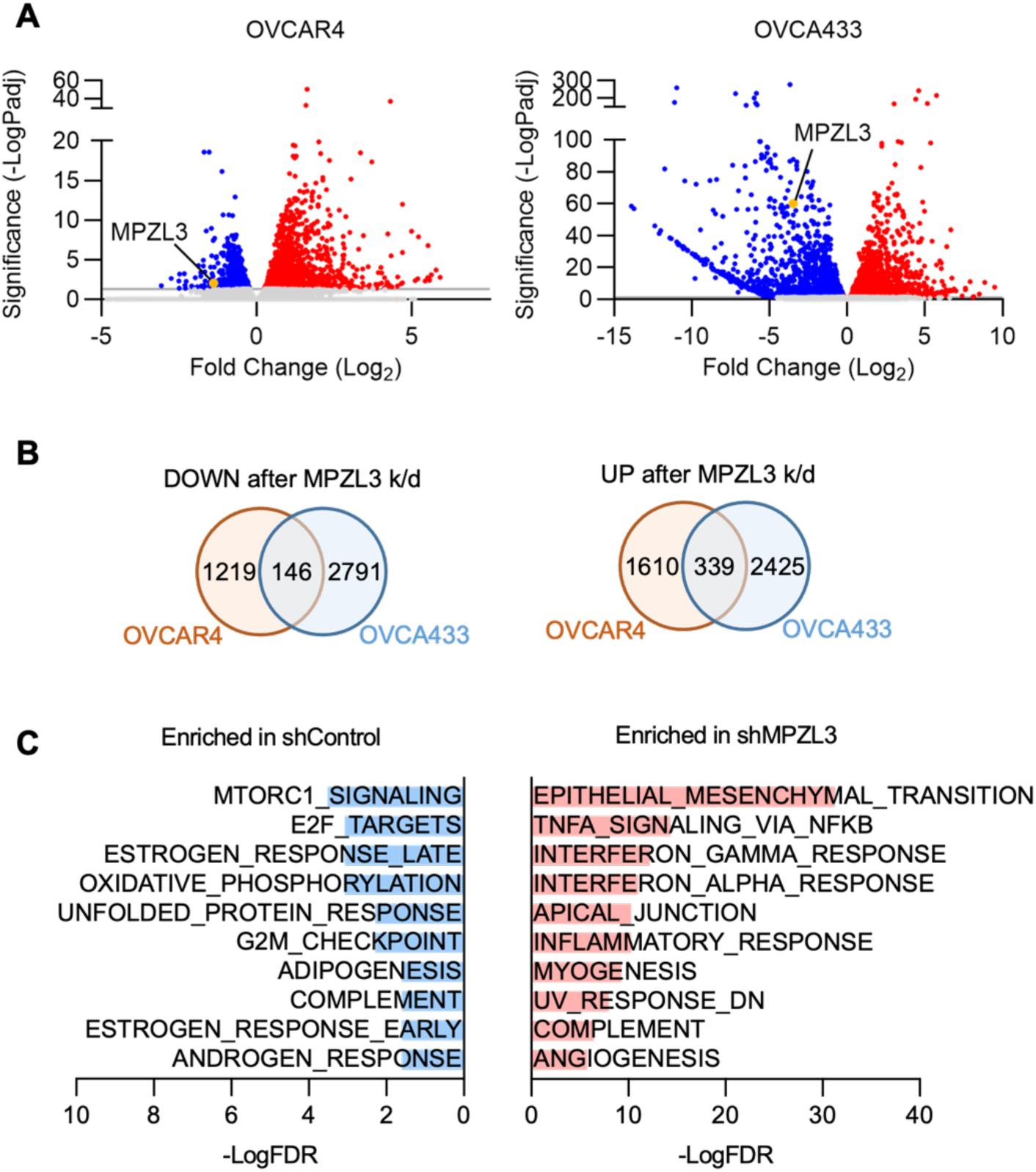
RNAseq analysis following MPZL3 knockdown. A. Distribution of selected DEGs in OVCAR4 and OVCA433 cells following MPZL3 knockdown (threshold: adjusted p-value < 0.05, log2 fold-change ≥ 1). B. Venn diagram of the number of significantly down and upregulated DEGs in OVCA433 and OVCAR4 cells following MPZL3 knockdown. C. Enriched Hallmark pathways commonly altered in OVCA433 and OVCAR4 cells following MPZL3 knockdown (MSigDB analysis of shared DEGs).

The role of CAMs in regulating EMT is well established [6, 35], but this has not been previously associated with MPZL3 loss. We found that the shared EMT signature genes set connected to MPZL3 knock-down in OVCAR4 and OVC433 cells (Figure 3A) was associated with significantly worse overall patient outcomes (TCGA Pan-Cancer; Figure 3B). Vimentin (VIM) and N-cadherin (CDH2) expression were further assessed at the protein levels in OVCAR4 cells following MPZL3 knockdown, which demonstrated a strong induction of vimentin expression in response to MPZL3 loss (Figure 3C&D), similar to that observed at the transcript level (Figure 3A & Supplementary Figure S2D). Moreover, vimentin expression was inversely correlated to MPZL3 expression in TCGA ovarian cancer specimens (Figure 3E), indicating that MPZL3 loss is associated with upregulation of the canonical EMT gene vimentin in ovarian cancer.

**Figure 3.**
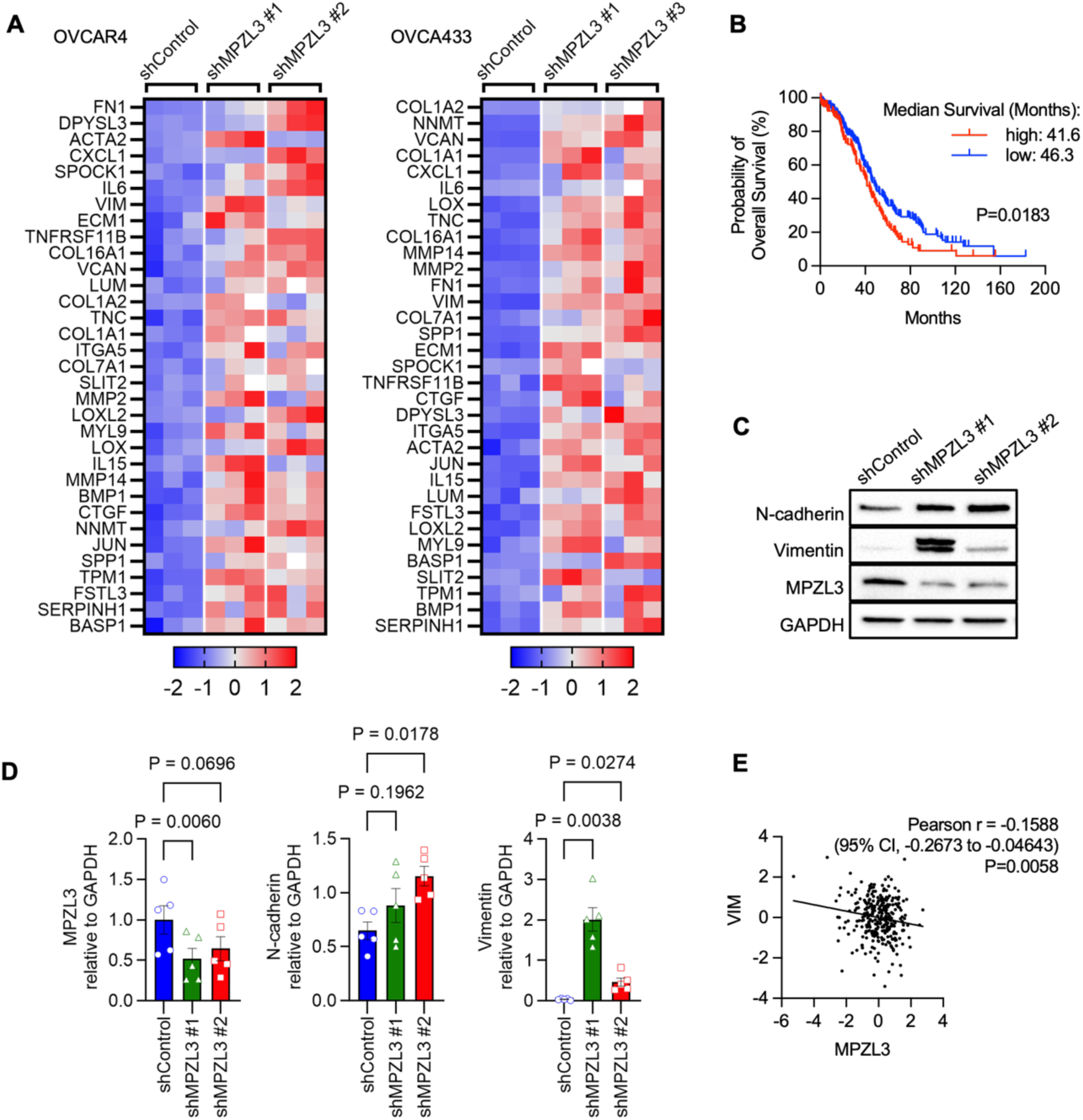
MPZL3 knock-down increases EMT gene expression signature which is associated with poor patent outcome. A. Heatmap of EMT signature gene transcripts increased in response to MPZL3 knock-down with two independent shRNAs in OVCAR4 and OVCA433 cells (RNA-seq, z-scores). B. Overall patient survival decreases with high expression of the MPZL3-dependent EMT gene signature (TCGA ovarian serous adenocarcinoma, Logrank Mantel-Cox test). C. Expression of Vimentin and N-cadherin protein levels are increased following MPZL3 knockdown in OVCAR4 cells. D. Densitometry quantification of C (n=5; one-way ANOVA, Dunnett’s test). E. Vimentin expression is negatively correlated to MPZL3 expression in TCGA ovarian serous adenocarcinoma specimens (mRNA z-scores; RNA Seq V2 RSEM).

### Loss of MPZL3 reduces homotypic cell adhesion and increases cell invasiveness in ovarian cancer

To determine if MPZL3 has CAM properties in ovarian cancer cells, we tested the effects of MPZL3 knockdown in OVCAR4 cells on homotypic cell adhesion by quantifying their attachment to a monolayer of parental OVCAR4 cells, and by assessing their heterotypic adhesion to mesothelial cells. Mesothelial cells line the abdominal cavity and represent the first barrier of attachment and invasion ovarian cancer cells face during metastasis in the peritoneal cavity. Loss of MPZL3 decreased homotypic adhesion between ovarian cancer cells but did not affect heterotypic cell adhesion to mesothelial cells (Figure 4A&B), suggesting that MPZL3 shares functional similarities to junctional adhesion molecules such as VSIG1. Finally, we conducted a mesothelial clearance assay [36], which demonstrated that MPZL3 knockdown promotes invasion and clearance of mesothelial cells (Figure 4C). In summary, our findings demonstrate that loss of MPZL3 induces EMT, reduces homotypic cell adhesion, and promotes invasion of ovarian cancer cells.

**Figure 4.**
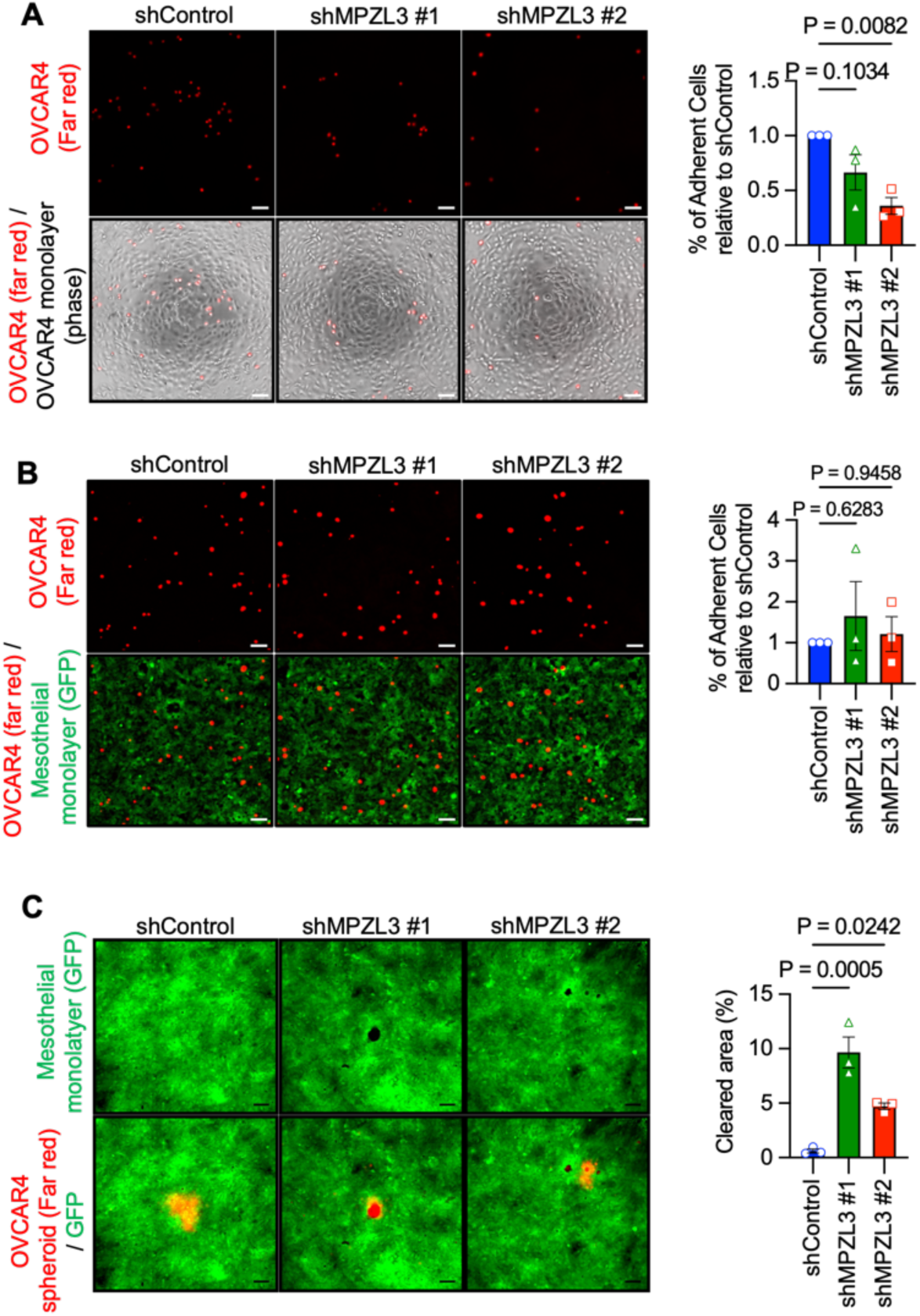
Loss of MPZL3 reduces homotypic cell adhesion and induces invasion of ovarian cancer cells. A. Homotypic adhesion was assessed by seeding parental OVCAR4 cells in a monolayer (phase contrast image) and quantifying adherence of OVCAR4 MPZL3 knockdown cells labeled with far-red fluorescent relative to the adherence of scramble control cells (scale bar: 100 µm; n=3; one-way ANOVA, Dunnett’s test). B. Heterotypic cell adhesion was assessed as in A, except OVCAR4 scramble control or MPZL3 knockdown cells labeled with far-red fluorescent dye were placed on GFP-labeled mesothelial cells (n=3; one-way ANOVA, Dunnett’s test). C. GFP-labeled mesothelial cells were seeded in a monolayer and mesothelial clearance by OVCAR4 scramble control or MPZL3 knockdown spheroids labeled with far-red fluorescent dye quantified after 48 hrs (scale bar: 100 µm; n=3; one-way ANOVA, Dunnett’s test).

### MPZL3 knockdown leads to cell cycle arrest and senescence

In addition to driving an EMT and invasive phenotype, MPZL3 knock-down led to a noticeable decrease in proliferation (Figure 5A, Supplementary Figure S3A). This correlated with a decrease in transcript expression associated with cell cycle genes (Figure 2C, Supplementary Figure S2C and S3B), including cyclin D1 (CCND1, Figure 5B). CCND1 and MPZL3 transcript expression were also significantly correlated in TCGA patient specimens (Figure 5C) and knock-down of MPZL3 resulted in a slowing in G0/1 to S/G2/M transition (Figure 5D, Supplementary Figure S3C) These findings suggest that loss of MPZL3 in ovarian cancer cells leads to G1 cell cycle arrest. We also observed that MPZL3 knock-down in OVCAR4 and OVCA433 cells led to a greater percentage of cells adopting a senescent-like phenotype, visualized using a β-galactosidase activity assay (Figure 5E and S3D), and this was accompanied by upregulation of common genes associated with the senescence-associated secretory phenotype (SASP) [37–42], such as *IL6* and *IL1A*, while *LMNB1* loss, another marker of senescence, was also observed (Figure 5F). We note that a population of these cells is still proliferating, which may be due to heterogenous knockdown in the culture.

**Figure 5.**
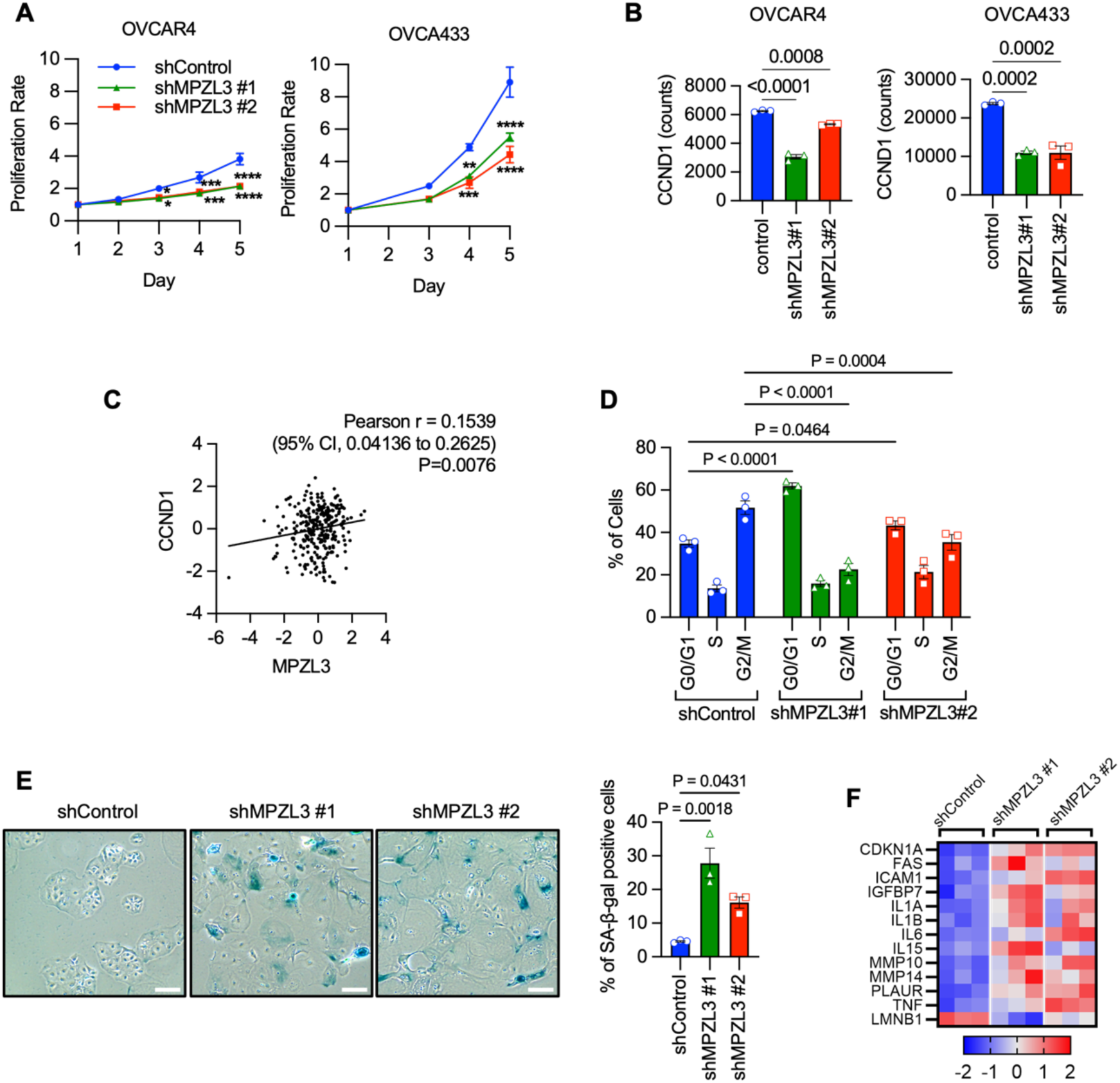
MPZL3 knockdown leads to cell cycle arrest and senescence. A. Cell growth assay following knock-down of MPZL3 in OVCAR4 and OVCA433 cells. Cell growth (FluoReporter Blue Fluorometric dsDNA Quantitation) was normalized to day 1 (n=3; two-way ANOVA, Dunnett’s test; *: P<0.05, **: P<0.01, ***: P<0.001, ****: P<0.0001). B. Cyclin D1 (CCND1) expression in OVCAR4 and OVCA433 cells following MPZL3 knock-down (RNASeq; n=3, one way ANOVA, Dunnett’s multiple comparisons test). C. Cyclin D1 (CCND1) expression positively correlates with MPZL3 expression in TCGA ovarian serous adenocarcinoma specimens (mRNA z-scores; RNA Seq V2 RSEM). D. Cell cycle analysis (PI Flow cytometry) of OVCAR4 cells following MPZL3 knock-down, at 9-hr time point from release (n=3; two-way ANOVA, Dunnett’s test). E. Images of senescence-associated (SA) β-galactosidase staining in OVCAR4 cells (scale bar: 200 µm). The corresponding quantification data are shown in the right panel (n=3; one-way ANOVA, Dunnett’s test). F. Heatmap of SASP markers and lamin B1 (LMNB1) following MPZL3 knockdown (OVCAR4 RNA-seq, z-scores).

### Loss of MPZL3 decreases sensitivity to cisplatin and olaparib

Given the observed slowing in proliferation, we hypothesized that loss of MPZL3 might increase drug resistance to chemotherapeutic agents. We determined the IC_50_ of three standard-of-care compounds used to treat ovarian cancer and observed that MPZL3 knock-down promoted resistance to Cisplatin and Olaparib, while no significant difference was observed with Paclitaxel treatment (Figure 6A and B). These data suggest that MPZL3 depletion decreases drug sensitivity, especially in response to treatments that induce DNA damage [43, 44]. To determine whether the increased Cisplatin resistance observed in MPZL3-depleted cells is recapitulated *in vivo*, we subcutaneously injected OVCAR4 cells expressing scramble control or MPZL3 shRNA#1 into NSG mice (Figure 6C-F). Once the average tumor volume in each group reached approximately 50 mm³, the mice were treated with either vehicle (saline) or Cisplatin. As expected from cell culture assays (Figure 5), MPZL3 knockdown tumors grew slower than scramble controls (Figure 6D&E). Comparing tumor growth rates in response to cisplatin treatment, the scramble control group significantly decreased their growth rate in response to Cisplatin treatment, while a significant change was not observed following MPZL3 knockdown (Figure 6E).

**Figure 6.**
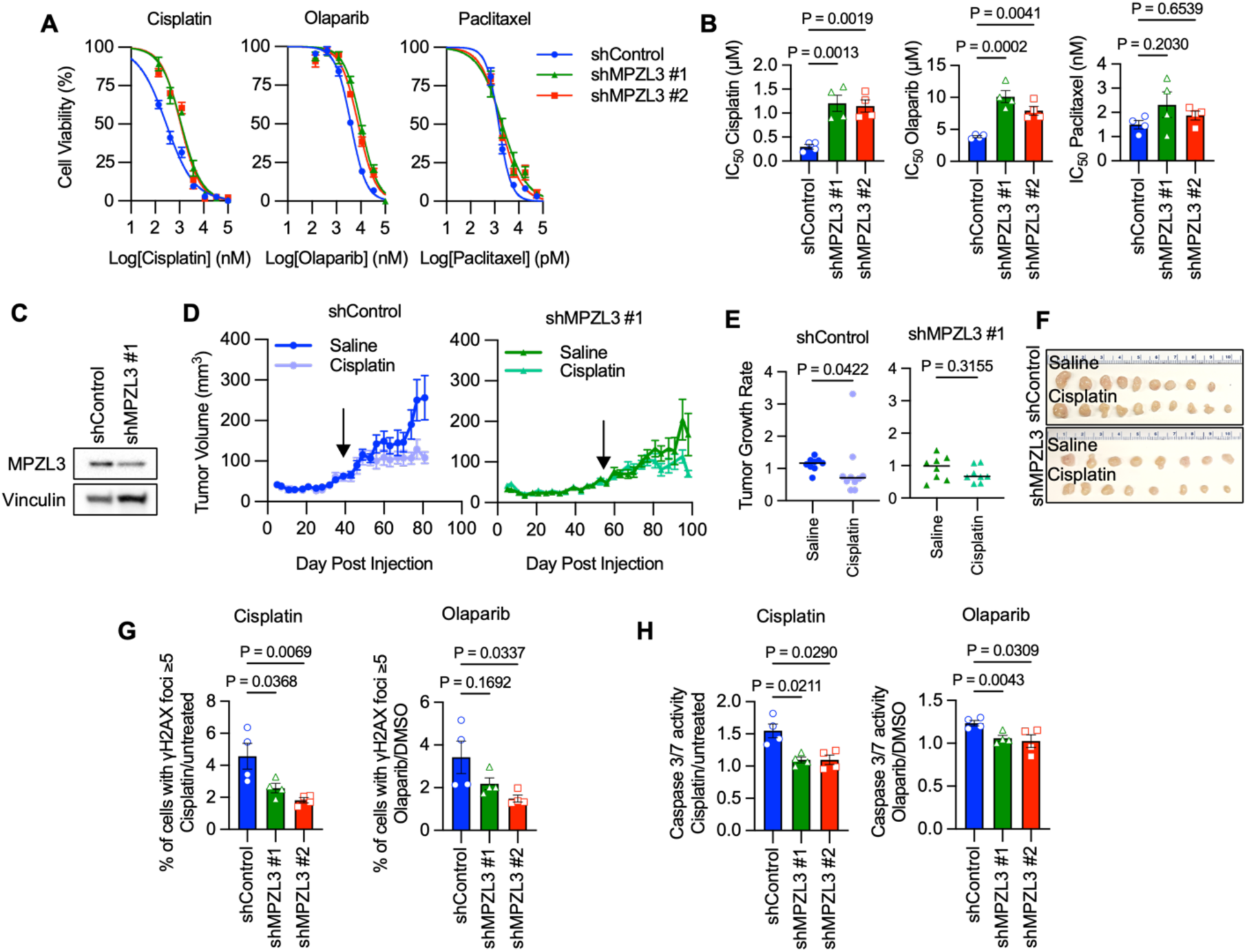
MPZL3 loss enhances Cisplatin and Olaparib resistance. A. Dose-response curves were derived from cell viability assays (FluoReporter dsDNA quantification) of OVCAR4 cells expressing scramble control or MPZL3 shRNAs (n=4, mean +/-SEM). B. IC_50_ values derived from dose-response curves in A (n=4; one-way ANOVA, Dunnett’s test). C. Western blot of MPZL3 knockdown OVCAR4 cells that were injected subcutaneously into NSG mice. D. Subcutaneous tumor growth of OVCAR4 cells expressing scramble control (left) or shMPZL3#1 (right) under vehicle (saline) or Cisplatin treatment (two tumors per mouse, n=4–5 mice per group). The start of Cisplatin treatment is indicated with an arrow. E. Tumor growth rates under treatments for both scramble control and shMPZL3#1 groups. Growth rates were calculated by dividing the final tumor volumes by the average tumor volumes from three measurements: three days before treatment, on the day treatment began, and four days after treatment began (median tumor volume shown, n=8-10, Mann-Whitney test). F. Final tumor volume images of OVCAR4 subcutaneous tumors from both saline- and Cisplatin-treated groups. All mice were euthanized after receiving the same doses of treatment. G. γH2AX signals induced by Cisplatin (left) or Olaparib (right) in OVCAR4 cells following MPZL3 knockdown (n=4; one-way ANOVA, Dunnett’s test). H. Caspase 3/7 activity induced by Cisplatin (left) or Olaparib (right) in OVCAR4 cells following MPZL3 knockdown (n=4; repeated measures one-way ANOVA, Dunnett’s test).

To further investigate how MPZL3 knockdown causes resistance to Cisplatin and Olaparib, γH2AX foci were assessed by immunofluorescence to monitor DNA damage response. Both Cisplatin and Olaparib treatments led to a higher percentage of cells with multiple γH2AX foci in scramble control cells compared to MPZL3 knockdown cells, indicating that MPZL3-depleted cells accumulate less DNA damage in response to Cisplatin or Olaparib stimulation. Considering that severe DNA damage response leads to apoptosis [45, 46], we assessed whether Cisplatin or Olaparib treatment induces more cell death in control cells compared to MPZL3-depleted cells. Cisplatin and Olaparib treatments induced higher levels of Caspase 3/7 activity in control cells compared to MPZL3 knockdown cells. This demonstrates that Cisplatin/Olaparib treatment causes less DNA damage and apoptosis in MPZL3-deficient cells and thus makes these cells less responsive to Cisplatin and Olaparib treatment.

## Discussion

In this study, we investigated the role of the novel CAM, MPZL3, in ovarian cancer. CNA analysis revealed that the MPZL3 locus, 11q23.3, is frequently altered across multiple cancer types (Figure 1). While the chromosome 11q23 region has been reported to be lost in many cancers [18–21], suggesting a tumor suppressor function for MPZL3, previously published bioinformatics data has suggested that MPZL3 is a prognostic marker in breast cancer [22]. Interestingly, we found a similar percentage of chromosome gains and losses at the MPZL3 locus in the TCGA ovarian adenocarcinoma dataset, leading us to question how MPZL3 influences ovarian cancer development and progression. Interestingly, we observed inter-patient heterogeneity related to *MPZL3* locus gain and loss, and it remains to be determined how this is reflective of MPZL3 expression levels during different stages of ovarian cancer initiation, progression, and metastasis. From our unbiased RNA-seq analysis performed on two ovarian cancer cell lines following MPZL3 knockdown, we observed that EMT gene sets were the most significantly enriched in the knockdown group and that cell proliferation and cell cycle gene sets were predominant in the control group (Figure 2). To functionally validate our findings, we confirmed that MPZL3 depletion reduced cell-cell adhesion between the same cell type, which for the first time demonstrates that MPZL3 acts as a CAM in epithelial cells, and that loss of this CAM drives an EMT phenotype in cancer cells that leads to enhanced mesothelial clearance and invasion (Figure 4). The finding that MPZL3 expression is further decreased in omental metastatic lesions suggests a potential role for MPZL3 loss during ovarian cancer metastasis (Figure 1D). Multiple CAMs have been shown to drive cancer progression by regulating cell-cell and cell-matrix interactions or acting as receptors for downstream signaling, ultimately influencing tumor growth and metastasis [35, 47, 48]. Both the loss and gain of specific CAMs are important features of EMT and metastasis. Our data suggests that loss of MPZL3 is reminiscent of the changes in E-cadherin observed during EMT and cancer metastasis.

Our findings demonstrating that MPZL3 loss drives both EMT and decreased proliferation are not surprising since a negative association between EMT and cell cycle progression has been reported [49]. Moreover, the connection between senescence, EMT, and drug resistance has also previously been established [42, 50–53]. Our data revealed that MPZL3 loss induces some cells to enter a senescent-like state, increases expression of SASP genes (Figure 5). In addition, we found increased resistance to Cisplatin and Olaparib, two compounds that elicit DNA damage or target DNA repair (Figure 6). Resistance to these compounds in MPZL3-depleted cells appears to be due to their decreased proliferation, resulting in reduced DNA damage accumulation and thus decreased apoptosis. This was somewhat surprising as senescence is generally associated with increased DNA damage, yet we did not see a consistent increase in DNA damage in response to MPZL3 knock-down alone in the absence of these compounds. Given that CAMs are implicated in both DNA damage response and drug resistance [54, 55], how MPZL3 loss may enhance DNA repair capacity requires further investigation.

MPZL3 contains an immunoglobulin-like domain that regulates immune cell recruitment [56, 57], indicating its potential role in mediating interactions between cancer cells and immune cells. However, the specific roles and mechanisms of MPZL3 in the TME, particularly in tumor immune infiltration and drug susceptibility in ovarian cancer, have not yet been studied. We analyzed the correlation between MPZL3 expression and immune infiltration levels using TIMER2.0 [58–60] and found that MPZL3 is positively correlated with CD8+ T cells, cancer-associated fibroblasts, and neutrophils, while negatively correlated with B cells in ovarian cancer (data not shown). Thus, it is possible that a loss of MPZL3 could also influence the recruitment of anti-tumor immune cells. In addition, we and others have previously implicated MPZL3 in the regulation of adipogenesis *in vivo* [8, 12]. Further studies are needed to determine if loss of MPZL3 similarly affects lipid metabolism in ovarian cancer cells. Notably, our RNA-seq data shows that the oxidative phosphorylation and adipogenesis gene set were enriched in control cells. We previously found that MPZL3 antisense knock-down can protect mice from diet-induced obesity and decrease adiposity and circulating lipids [8]. Interestingly, one of the target genes of MPZL3 loss, vimentin, has been implicated in lipolysis, lipid uptake, and lipid droplet transport [61–63]. Although *VIM*-/- mice appear to display a similar metabolic phenotype compared to MPZL3-/- mice [64, 65]. Nevertheless, whether changes in vimentin also drive altered lipid metabolism in ovarian cancer cells is an intriguing hypothesis, particularly since lipid metabolism is a phenotype of tumor cells metastasizing to the omentum.

These data demonstrate for the first time that loss of the CAM MPZL3 results in EMT, decreases proliferation, and drug resistance in ovarian cancer, and that decreased MPZL3 expression is a phenotype of ovarian cancer tumor metastasis and may contribute to treatment failure in advanced-stage disease patients.

## Acknowledgements

The authors thank Weihua Pan, Sara Shimko and Ben Yankasky for technical assistance. We thank Dr. Yannis Zervantonakis for kindly providing mesothelial cells. This work was supported by NIH R01CA242021 (N.H.), and NIH training grant 2T32HL110849-11A1 (to S.W.). Lauren Borho and Dr. Francesmary Mudugno kindly assisted as honest brokers to access patient specimens. The ProMark tissue bank is supported by NIH SPORE P50CA272218. This project used the Hillman Cancer Center Cytometry Facility, Animal Facility, In vivo Imaging Facility and Cell and Tissue Imaging Facility that are supported in part by award P30CA047904.

