## Supplemental Figures for "Loss of the predicted cell adhesion molecule MPZL3 promotes EMT and chemoresistance in ovarian cancer"

**Cheng et al. 2024**

### Supplementary Figures

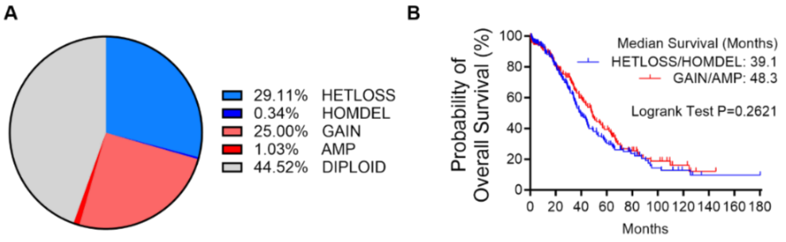


**Supplementary Figure S1. Percentage of *MPZL3* CNA and CNA based overall survival in TCGA ovarian adenocarcinoma dataset.**

A. Percentage of various MPZL3 CNA in ovarian adenocarcinoma samples (TCGA, PanCancer Atlas).
B. Overall patient survival in relation to *MPZL3* copy number. Loss of MPZL3 is negatively associated with patient survival (Logrank Mantel-Cox test).

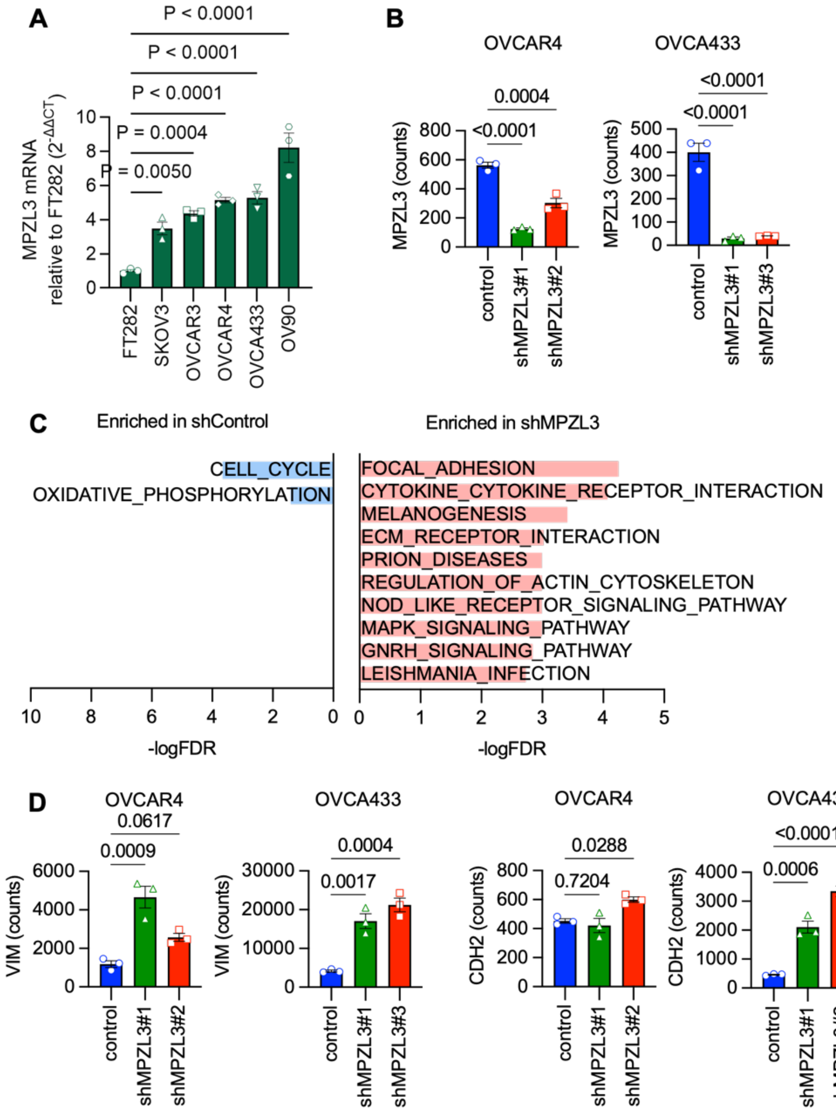


**Supplementary Figure S2. MPZL3 knock-down affects transcription of EMT and cell cycle-related genes in ovarian cancer cells.**

A. *MPZL3* mRNA expression in ovarian cancer cell lines (n=3; one-way ANOVA, Dunnett's test)
B. MPZL3 expression following shRNA mediated knock-down in OVCAR4 and OVCA433 (counts from RNA sequencing analysis).
C. Enriched KEGG pathways commonly altered in OVCA433 and OVCAR4 cells following MPZL3 knockdown (MSigDB analysis of shared DEGs).
D. Vimentin (VIM) and N-cadherin (CDH2) expression in OVCAR4 and OVCA433 cells following MPZL3 knock-down (RNASeq; n=3, one way ANOVA, Dunnett's multiple comparisons test).

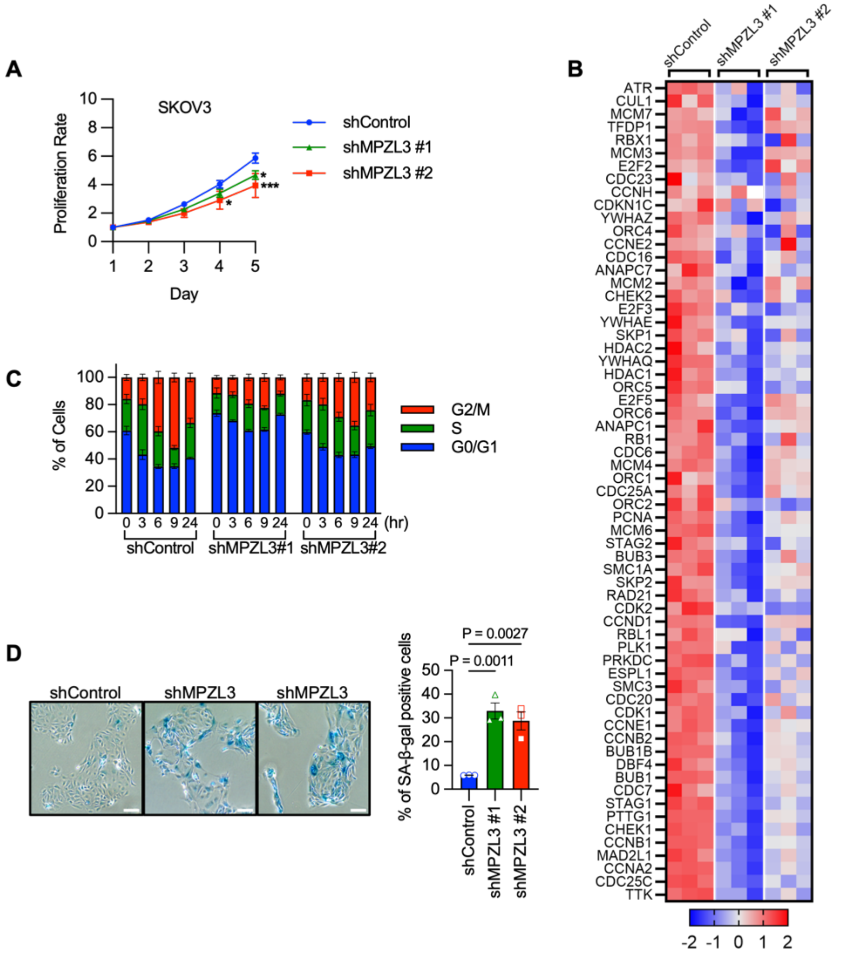


**Supplementary Figure S3. MPZL3 knock-down decreases proliferation and cell cycle progression**

A. Cell growth assay results for SKOV3 cells following MPZL3 knock-down. Results were normalized to day 1 to obtain the normalized ratio (N=3; two-way ANOVA, Dunnett's test; *: $P<0.05$, **: $P<0.01$, ***: $P<0.001$, ****: $P<0.0001$).
B. Heatmap of core enrichment genes from the cell cycle gene set following MPZL3 knockdown (KEGG, OVCAR4 RNA-seq, z-scores).
C. PI Flow cytometry analysis in OVCAR4 cells with MPZL3 knockdown. Cells were synchronized at the G1/S phase, released and harvested at 0, 3, 6, 9, and 24 hrs (n=3).
D. Images of senescence-associated (SA) β-galactosidase staining in OVCA433 cells (scale bar: 200 μm). The corresponding quantification data are shown in the right panel. (N=3; one-way ANOVA, Dunnett's test).
